# DNA Nanostructures Coordinate Gene Silencing in Mature Plants

**DOI:** 10.1101/538678

**Authors:** Huan Zhang, Gozde S. Demirer, Honglu Zhang, Tianzheng Ye, Natalie S. Goh, Abhishek J. Aditham, Francis J. Cunningham, Chunhai Fan, Markita P. Landry

**Author notes:** Authors contributed equally to this work (Huan Zhang, Gozde S. Demirer, Honglu Zhang). **Significance** Plant bioengineering will be necessary to sustain plant biology and agriculture, where the delivery of biomolecules such as DNA, RNA, or proteins to plant cells is at the crux of plant biotechnology. Here, we show that DNA nanostructures can passively internalize into plant cells and deliver small interfering RNA (siRNA) to mature plant tissues. Furthermore, we demonstrate that nanostructure size, shape, compactness, and stiffness, affect both nanostructure internalization into plant cells and subsequent gene silencing efficiency. Interestingly, we also find that the siRNA attachment locus affects the endogenous plant gene silencing pathway. Our work demonstrates programmable passive delivery of biomolecules to plants, and details the figures of merit for future implementation of DNA nanostructures in agriculture.

## Abstract

Plant bioengineering may generate high yielding and stress-resistant crops amidst a changing climate and a growing global population (1–3). However, delivery of biomolecules to plants relies on *Agrobacterium* infection (4) or biolistic particle delivery (5), the former of which is only amenable to DNA delivery. The difficulty in delivering functional biomolecules such as RNA to plant cells is due to the plant cell wall which is absent in mammalian cells and poses the dominant physical barrier to exogenous biomolecule delivery in plants. DNA nanostructure-mediated biomolecule delivery is an effective strategy to deliver cargoes across the lipid bilayer of mammalian cells, however, nanoparticle-mediated delivery remains unexplored for passive biomolecule delivery across the cell wall in plants. Herein, we report a systematic assessment of different DNA nanostructures for their ability to internalize into cells of mature plants, deliver small interfering RNAs (siRNAs), and effectively silence a constitutively-expressed gene in *Nicotiana benthamiana* leaves. We show that nanostructure internalization into plant cells and the corresponding gene silencing efficiency depends on the DNA nanostructure size, shape, compactness, stiffness, and location of the siRNA attachment locus on the nanostructure. We further confirm that the internalization efficiency of DNA nanostructures correlates with their respective gene silencing efficiencies, but that the endogenous gene silencing pathway depends on the siRNA attachment locus. Our work establishes the feasibility of biomolecule delivery to plants with DNA nanostructures, and details both the design parameters of importance for plant cell internalization, and also assesses the impact of DNA nanostructure geometry for gene silencing mechanisms.

## Introduction

Plant bioengineering has great potential to generate high yielding and pathogen-resistant crops, and is at the core of sustainability efforts (1, 2). However, unlike mammalian cells, plant cells have a cell wall which poses the dominant barrier to exogenous biomolecule delivery. Currently, biological delivery (using bacteria or viruses) and particle bombardment are the two preferred methods of biomolecule delivery to plant cells. However, biological delivery methods are highly cargo and host-specific (6) whereas particle bombardment can result in tissue damage (7). Nanomaterial-mediated biomolecule delivery has facilitated genetic engineering and biosynthetic pathway mapping in animal systems (8–12), but has only recently been explored for plants. Specifically, two recent studies have shown that carbon nanotubes (13) and clay nanosheets (14) enable passive intracellular delivery of DNA and RNA through surface-grafting or encapsulation strategies, circumventing the use of biolistics (external force). Passive biological cargo delivery to plants is an exciting development that warrants an understanding of how nanomaterials can passively internalize into plant cells, so that nanotechnology can be logically designed for future applications in plant biotechnology.

DNA nanotechnology leverages the programmability of DNA Watson-Crick base pairing to assemble DNA nanostructures into custom predesigned shapes via sequence-specific hybridization of template and staple DNA strands (15). To date, a plethora of different DNA nanostructures of variable sizes and shapes have been synthesized (16–19), and have shown functionality in biotechnology (20) for drug, DNA, RNA, and protein delivery applications in animal systems (21–26). However, to-date, DNA nanostructures have not been explored for use in plant systems, despite their utility shown in other sectors of biotechnology.

Herein, we explore DNA nanotechnology as a biomolecule delivery platform in plants. We synthesized a series of DNA nanostructures of controllable size, shape, stiffness, and compactness, and designed attachment loci onto which DNA, RNA, or protein cargoes may be conjugated. By hybridizing fluorophore-conjugated DNA strands onto the loci of DNA nanostructures, we tracked nanostructure internalization into the plant cell cytoplasm and found that stiffness and size are important design elements for nanostructure internalization into plant cells. DNA nanostructures with sizes below 10 nm, and higher stiffness or compactness, showed higher cellular internalization – although size or stiffness alone are not mutually-exclusive contributors to nanostructure internalization. DNA nanostructures were next loaded with siRNA targeting a GFP gene and infiltrated into plant leaves. This study revealed that DNA nanostructures enable gene silencing in plant leaves with efficiencies that match nanostructure internalization trends. Interestingly, the plant endogenous gene silencing mechanism can be affected by the DNA nanostructure shape and the siRNA attachment locus, affecting whether silencing occurs dominantly through transcriptional or post-transcriptional gene silencing. Our study confirms that DNA nanostructures can be designed to passively internalize into plant cells, and that DNA nanostructures may be a promising toolset for the delivery of exogenous biomolecules to plants, as has proven valuable in animal systems.

## Results

### Design, construction, and characterization of DNA nanostructures

We report the synthesis and systematic assessment of different DNA nanostructures for their ability to internalize passively into plant cells, and their subsequent utility for the delivery of siRNAs to mature plants. Three DNA nanostructures with programmed sizes and shapes were synthesized: a zero-dimensional (0D) **tetrahedron**, a one-dimensional (1D) hairpin-tile (**HT**) monomer, and a high-aspect-ratio 1D **nanostring**(27) as illustrated in Figure 1 (methods and Table S1). Both the HT monomer and tetrahedron were assembled through four single stranded DNA oligonucleotides. Briefly, the HT monomer structure was designed to contain a sticky end and a stem-loop hairpin structure, enabling co-polymerization with another monomer to assemble into the length-controlled 1D nanostring by introduction of an initiator (supplementary methods). The tetrahedron was also assembled through annealing of four pre-designed single stranded DNA oligonucleotides. Based on B-form double helix DNA dimensions (2 nm diameter, 0.33 nm per base in the direction of the helical axis), the sizes of the nanostructures are 2 × 5 × 16 nm for the HT monomer; 2 × 5 × 320 nm for the 10-unit nanostring; and 2.4 nm for all edges of the tetrahedron. Atomic force microscopy (AFM) characterization in Figure S1 and S2 shows proper formation of the DNA nanostructures.

Each nanostructure was programmed to attach a biological cargo – DNA, RNA, or protein – to a predefined locus or loci through complementary base pair hybridization. As visualized in Figure 1, the tetrahedron contained one attachment locus at its apex, the nanostring contained ten attachment loci at the center of each of its constituent monomers, and the HT monomer contained one attachment locus either at its center (HT-c), or for a separate construct, an attachment locus at its side (HT-s). To confirm the accessibility of the attachment loci, streptavidin protein was attached to the siRNA attachment locus (supplementary methods) to visualize the conjugation site in the HT monomer and nanostring. AFM imaging revealed the predicted attachment of one streptavidin protein in the center or side of the HT-c or HT-s monomers, respectively, and ten streptavidin proteins per nanostring at the center of each constituent HT monomer (Figure 1). Confirming the synthesis and attachment loci of the DNA nanostructures motivated an assessment of their passive internalization propensities into the cells of mature plant tissues.

**Figure 1.**
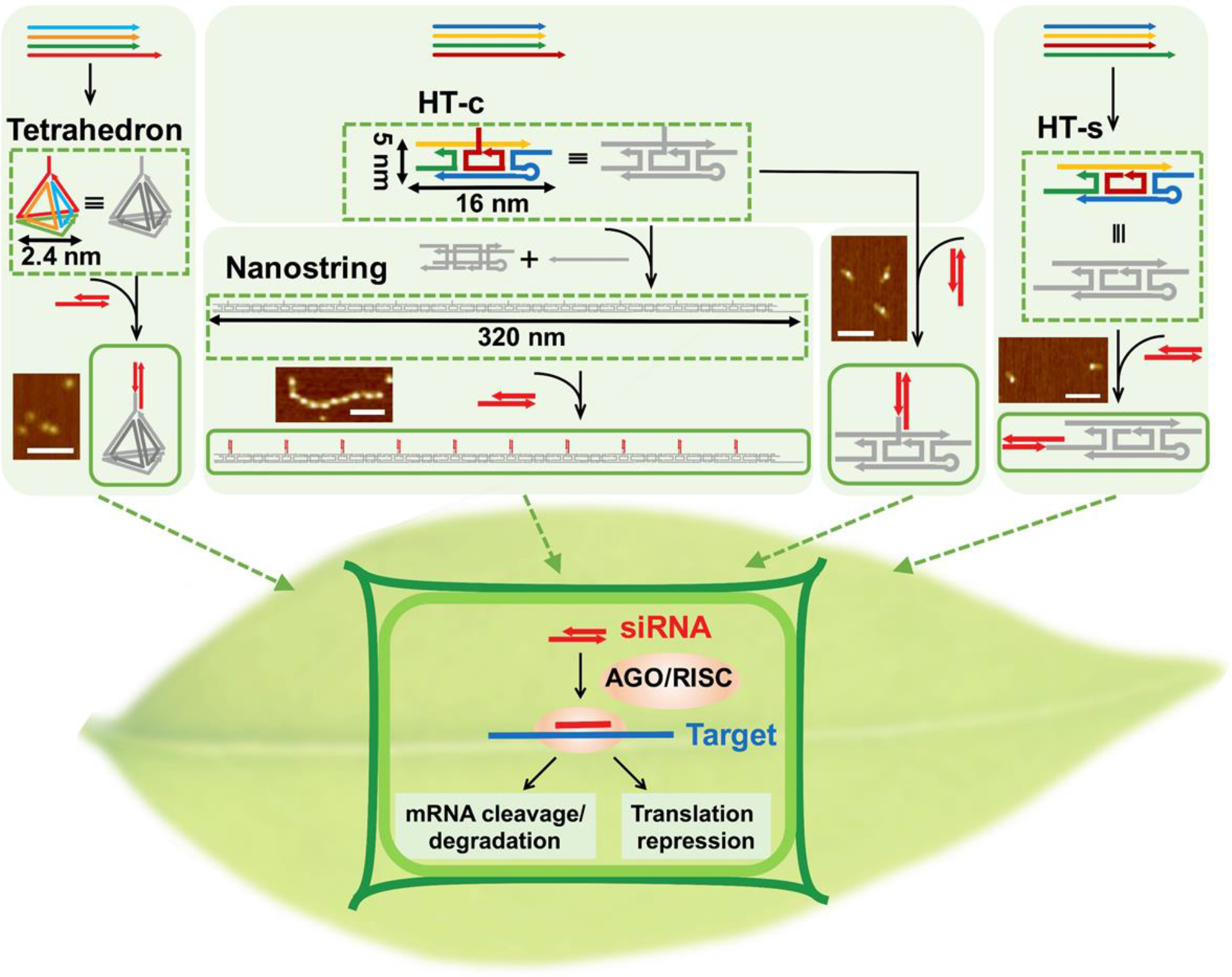
DNA nanostructure synthesis and plant infiltration workflow. Tetrahedron and hairpin-tile (HT) monomer nanostructures were synthesized from four single stranded DNA sequences, and the 1D nanostring structure was synthesized by polymerization of HT monomers. The cargo (DNA, RNA, protein) attachment locus was designed to be at the apex of the tetrahedron, along the length of the nanostring, and to the side (HT-s) or center (HT-c) of each hairpin-tile (HT) nanostructure. AFM imaging of streptavidin binding to the biotinylated nanostructure locus confirms proper attachment of cargo on the HT monomer and nanostring nanostructures. DNA nanostructures loaded with siRNA at each locus are infiltrated into the transgenic mGFP5 *Nicotiana benthamiana* plant leaves for downstream study of nanostructure internalization and gene silencing. Scale bars = 100 nm.

### Internalization mechanism of DNA nanostructures into plant cells

While the size exclusion limit set by the cell membrane is estimated to be around 500 nm, the plant cell wall has been reported to exclude particles larger than 5-20 nm (28). Motivated by this figure of merit, we tested whether DNA nanostructures could passively internalize into the cells of *Nicotiana benthamiana* leaves. DNA nanostructures were fluorescently labeled via attachment of Cy3 labeled DNA strands to nanostructure attachment loci (supplementary methods) and infiltrated into the leaf abaxial side to assess cellular uptake in mGFP5 *Nicotiana benthamiana* transgenic plants (Figure 2a). Confocal microscopy imaged both the Cy3 fluorescence of the nanostructures concurrently with the intrinsic cytosolic GFP fluorescence generated by the plant cells. Colocalization of the Cy3 fluorescence (nanostructure) with the GFP fluorescence (plant cell cytosol) 12-hours post-infiltration was used to determine the extent of nanostructure internalization into the cell cytosol. Colocalization analysis (supplementary methods) in Figure 2b shows that the HT monomer and tetrahedron nanostructures exhibit a high degree of colocalization with the plant cell cytosol (59.5 ± 1.5% and 54.4± 2.7 % mean ± SD, respectively), while the nanostring showed a lower degree of colocalization (35.8 ± 0.9% mean ± SD) with the plant cell cytoplasm. Representative confocal microscopy images of colocalization are shown in Figure 2c and Figure S3, suggesting HT monomers and tetrahedra passively internalize in to plant cells significantly less than nanostrings. We observe that a large portion of the Cy3 fluorescence from the Cy3-nanostring infiltrated leaves originates from nanostrings that are putatively stuck in the guard cells, which is the dominant contribution to the colocalization fraction calculated for nanostring internalization (Figure 2b, Figure S4). Conversely, we observe that most of the Cy3 fluorescence recovered from Cy3-HT infiltrated leaves follows the cell contour, identified by cytosolic GFP expression. Furthermore, free Cy3-oligonucleotides alone infiltrated into the leaves did not show significant colocalization with the cell cytoplasm (18.0 ± 4.6 % mean ± SD, Figure S5).

We next tested whether the cellular uptake mechanism is predominantly an energy dependent or independent process by infiltrating the Cy3-labeled HT monomer into mGFP5 *Nicotiana benthamiana* plant leaves at either 20°C or 4°C, where at 4°C, energy-dependent cellular uptake is reduced (29). As shown in Figure S6, most of the Cy3-labeled monomer nanostructure is retained around the leaf stomata (guard cells) upon infiltration and incubation at 4°C, whereas Cy3-labeled monomer nanostructures enter and diffuse uniformly into the cell cytoplasm if the infiltration and incubation is performed at 20°C. Therefore, we propose that HT monomer nanostructures are actively uptaken through the plant cell membrane by an energy-dependent mechanism.

**Figure 2.**
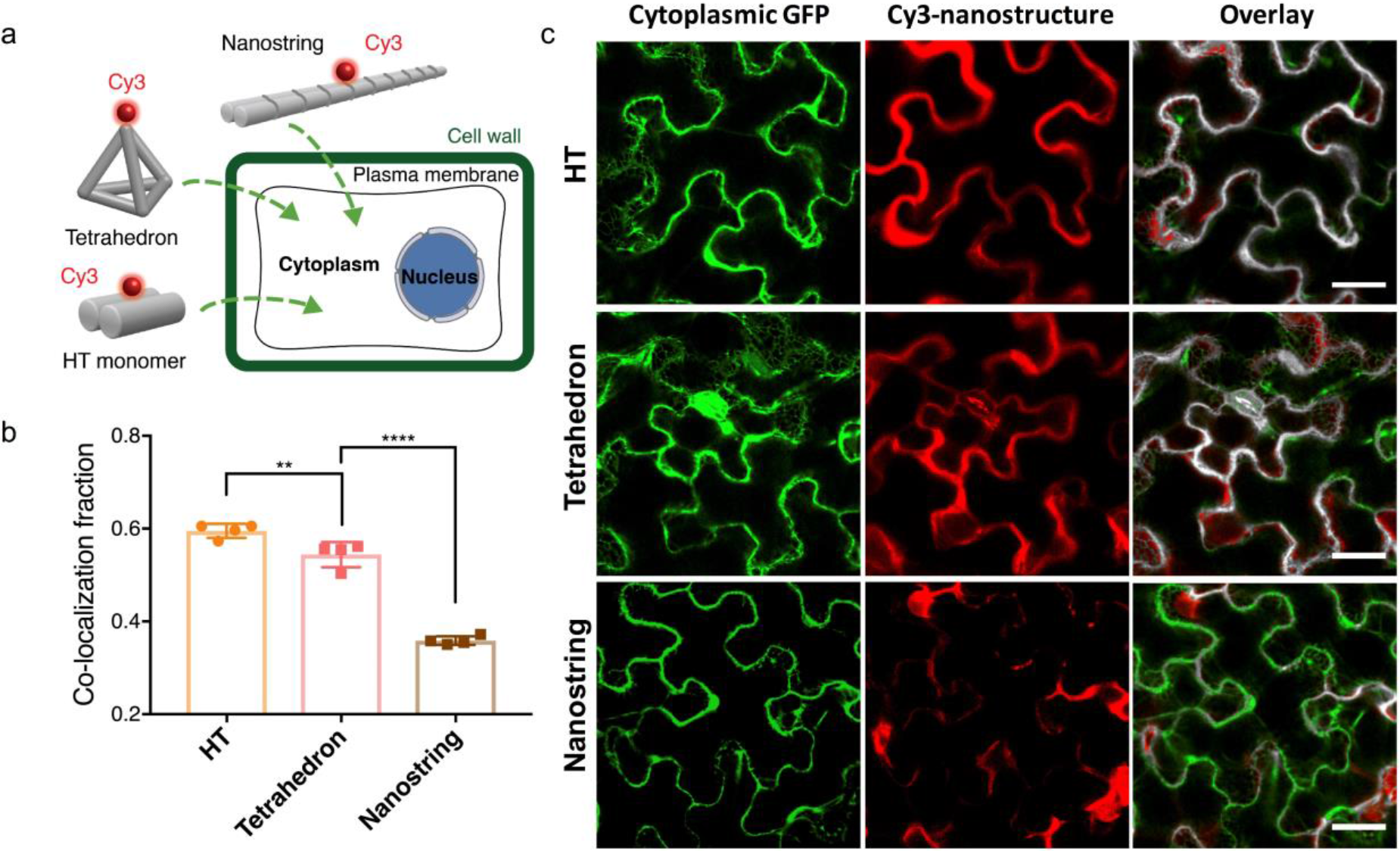
DNA nanostructure internalization into and colocalization with mGFP5 *Nicotiana benthamiana* cytoplasm. a) Internalization assays of Cy3-tagged DNA nanostructures into mGFP5 *benthamiana* plant cells. b) Colocalization of Cy3 fluorescence (nanostructure) with GFP fluorescence (plant cell cytosol) 12-hours post-infiltration into mGFP5 *benthamiana* leaves. P** = 0.0041 and P**** < 0.0001 in one-way ANOVA. Error bars indicate s.e.m. (n = 4). c) Representative confocal images from data in (b) for HT, tetrahedron, and nanostring DNA nanostructures.

Prior work probing nanomaterial uptake in mammalian systems suggests uptake across the lipid bilayer is dependent on nanoparticle size, shape, aspect ratio, and mechanical stiffness (30, 31). We posit these parameters may also affect DNA nanostructure uptake across the plant cell wall. To better understand nanostructure parameters enabling passive plant cell internalization, we compiled and compared the size, compactness, aspect ratio (after conjugation with siRNA), and relative stiffness of the DNA nanostructures. Regarding size, as shown in Table S2, we find that efficient nanostructure uptake into plant cells requires at least one dimension at or below ~ 5 nm, possibly to remain below the plant cell wall size exclusion limit (32, 33). Regarding compactness (supplementary methods and Table S2), we calculated that the tetrahedron and HT monomer exhibit higher compactness than the nanostring (0.55 and 0.45 vs 0.11, respectively). Our results indicate that nanostructures with higher compactness result in higher cellular uptake efficiency (59.5 ± 1.5 % for HT monomer, and 54.4± 2.7 % for tetrahedron, compared to 35.8 ± 0.9% for nanostring) consistent with mammalian cell systems (34).

To further rationalize DNA nanostructure internalization trends, we calculated and compared the relative bending stiffness (ratio of bending stiffness of dsDNA, to bending stiffness of the nanostructure kb _structure_/kb _dsDNA_, Table S2) of the HT monomer and nanostring. In comparing the HT monomer to the nanostring, we find that a higher nanostructure stiffness (0.83 vs. 1 × 10^−4^ kb_nanostructure_/kb_dsDNA_ for the HT monomer versus nanostring, respectively) results in higher cellular uptake (59.5 ± 1.5 % vs. 35.8 ± 0.9 % ; mean ± SD for the monomer versus nanostring, respectively). We also simulated the flexibility of siRNA alone, HT, and nanostring DNA nanostructures using CanDo (35, 36) (supplementary methods) to understand how nanostructure stiffness affects plant cell internalization. Mechanical stiffness calculations quantifying the root mean square fluctuations (RMSF), as summarized in Figure S7, demonstrate that the nanostring is significantly more flexible than the HT monomer (supplementary video 1 and 2). Based on the above calculations, we hypothesize that in addition to nanostructure size, the mechanical stiffness of the nanostructure plays an important role in nanostructure internalization into plant cells.

To further explore this hypothesis, we tested the effect of nanostructure stiffness on plant cell internalization. Single-walled carbon nanotubes (SWCNTs) exhibit stiffnesses on the order of hundreds of GPa to TPa (37, 38), about one thousand times stiffer than the nanostring nanostructure (several Gpa (39)). Furthermore, SWCNTs have been shown to passively internalize into the cells of a variety of plant species (40–42), with a leading hypothesis that the larger tensile strength of the SWCNT compared to that of the plant cell wall facilitates needle-like plant cell internalization (13, 43, 44). To test the effect of nanostructure stiffness for internalization into *Nicotiana benthamiana* plant cells, we tethered DNA nanostrings to SWCNTs (supplementary methods). AFM characterization and near-infrared spectroscopy confirmed successful nanostring-SWCNT conjugation (Figure S8; supplementary methods). As shown in Figure S9, colocalization analysis indicates that while the nanostring alone does not internalize into the plant cell, the nanostring-SWCNT conjugate does, with a similar internalization efficiency (51.4 ± 4.5 % mean ± SD) as the internalization efficiency of Cy3-labeled SWCNTs alone (54.2± 4.5 % mean ± SD).

These results suggest that tethering a flexible nanostring to the relatively inflexible SWCNT enables nanostring transport into the plant cell, despite the fact that the nanostring-SWCNT conjugate is larger in size than either the nanostring or the SWCNT alone. Based on these results, we conclude that the nanostructure stiffness is a key design element for nanostructure internalization into plant cells. In summary, we find that size (at least one dimension below ~5 nm), higher aspect ratio, and higher stiffness are all contributing figures of merit for nanostructure internalization into plant cells, as summarized in Table S2.

### siRNA-mediated gene silencing efficiency of DNA nanostructures

We next examined whether DNA nanostructures could be loaded with a functional biological cargo, siRNA, to accomplish gene silencing in plants. RNA interference (RNAi) is a phenomenon in which double stranded RNA (dsRNA) induces gene silencing, and has expedited discoveries in genomics and therapeutics (45). A key conserved feature of RNAi in plants is processing of dsRNA into siRNAs by the activity of Dicer-like enzymes (45, 46). siRNAs are subsequently incorporated into an RNA-induced silencing complex (RISC), resulting in sequence-specific blocking of mRNA translation. RNAi has emerged as a powerful strategy to engineer disease resistance against pests and pathogens in plants, and has facilitated plant biosynthetic pathway mapping.

To ascertain whether DNA nanostructures can deliver siRNA and achieve gene silencing in plants, we targeted the silencing of a GFP gene in transgenic mGFP5 *Nicotiana benthamiana*, which exhibit strong constitutive GFP expression from the nuclear genome. We designed a 21-bp siRNA sequence that inhibits GFP expression in a variety of monocot and dicot plants (47), and hybridized this duplex oligonucleotide to a complementary strand programmed into the site-specific loci on the DNA nanostructures (Table S1 for sequences). Native polyacrylamide gel electrophoresis (PAGE, Figure S10) or agarose gel electrophoresis analysis (Figure S11) was performed to validate conjugation of siRNA to each DNA nanostructure. Furthermore, we confirmed that DNA nanostructures remain stable in various biological media for at least 12 hours (Figure S12), motivating their use in plant tissues.

Following siRNA loading, each nanostructure with its linked active siRNA duplex(es) was introduced into the leaves of mGFP5 *Nicotiana Benthamiana* via infiltration to the leaf abaxial side for GFP gene silencing experiments, with an siRNA concentration of 100 nM (Figure 3a). Confocal microscopy was performed to image GFP expression in infiltrated leaves, and western blotting was utilized as a second method to confirm and quantify GFP expression changes in the infiltrated leaf tissues. As shown in representative confocal images in Figure 3b, untreated control leaves or leaves treated with free siRNA alone showed strong GFP fluorescence (low or no gene silencing), as expected, due to constitutive expression of GFP in the transgenic plant. Conversely, leaves infiltrated with siRNA-linked DNA nanostructures showed varying degrees of reduced GFP fluorescence 3-days post-infiltration.

Specifically, as shown in Figure 3c, leaves infiltrated with siRNA functionalized nanostrings showed a ~29 ± 4.6% (mean ± SD) decrease of GFP fluorescence compared to the untreated leaf. Leaves infiltrated with the HT monomer showed a 41 ± 5.4 % or 47 ± 4.7% (mean ± SD) reduction in GFP fluorescence for constructs in which the siRNA was linked at the center or side of the nanostructure, respectively. Lastly, leaves infiltrated with siRNA conjugated to the tetrahedron showed a 42% ± 6.5 %, (mean ± SD) decrease in GFP fluorescence intensity compared to untreated leaves (Figure 3c). Notably, all leaves infiltrated with siRNA functionalized DNA nanostructures exhibited a significantly larger fluorescence decrease compared to leaves infiltrated with free siRNA, suggesting that DNA nanostructures can serve as a nucleotide delivery tool in plant systems.

We note that the degree of nanostructure internalization (Figure 2) is proportional to the silencing efficiency achieved with each nanostructure (Figure 3), suggesting that nanostructure internalization into the plant cell determines its ability to induce siRNA-based gene silencing. Interestingly, we observe higher (47 ± 4.7 % ; mean ± SD) gene silencing efficiency when siRNA is linked to the side of the HT monomer (HT-s, aspect ratio 5:1), compared to a lower (41 ± 5.4 %; mean ± SD) silencing efficiency when siRNA is instead linked to the center of the HT monomer (HT-c, aspect ratio 1:1). These results are congruent with prior studies suggesting that higher aspect ratio nanostructures facilitate nanoparticle entry into cells (28). However, the nanostring, which has the highest aspect ratio (20:1), surprisingly shows the lowest silencing efficiency (29 ± 4.6%; mean ± SD) and internalization efficiency (35.8 ± 0.9 %; mean ± SD), suggesting that nanostructure shape is not the only parameter affecting internalization into plant cells. In particular, above internalization assays show nanostring internalization into plant cells *only* if the nanostring is first conjugated to a high stiffness nanostructure such as a SWCNT, confirming that nanostructure stiffness is an important parameter for both nanostructure internalization and gene silencing efficiency.

To further confirm siRNA-induced decrease in GFP, western blotting was applied to quantify GFP expression in each plant leaf 3 days post-infiltration with the siRNA-linked nanostructure (supplementary methods). As shown in Figure 3d, the HT monomer and tetrahedron nanostructures linked with siRNA show a significant decrease in GFP compared to untreated leaves: 37 ± 4.3 % GFP decrease for HT-c, 49 ± 3.8 % decrease for HT-s, and 40 ± 1.9% decrease for the tetrahedron (mean ± SD). Interestingly, the siRNA conjugated to the *end* of the HT monomer nanostructure showed the best silencing efficiency and the most GFP decrease, which was significantly higher than when the siRNA was instead conjugated to a locus on the *center* of the HT monomer. Moreover, we observed no statistically significant silencing by the siRNA loaded nanostring compared to siRNA alone.

We also tested the transience of the nanostructure-enabled siRNA mediated gene silencing. Confocal imaging shows that GFP fluorescence for all siRNA-loaded DNA nanostructure treated leaves recovers to pre-infiltration or non-infiltration (control) levels by 7-days post-infiltration (Figure S13). Transience of siRNA-mediated gene silencing was also verified by quantifying GFP expression with quantitative western blot analysis. As shown in Figure 3e, the amount of GFP expressed in the leaves infiltrated with the HT-s monomer and tetrahedron nanostructures, which had induced the largest GFP silencing at day 3, returned to baseline protein expression levels by day 7.

**Figure 3.**
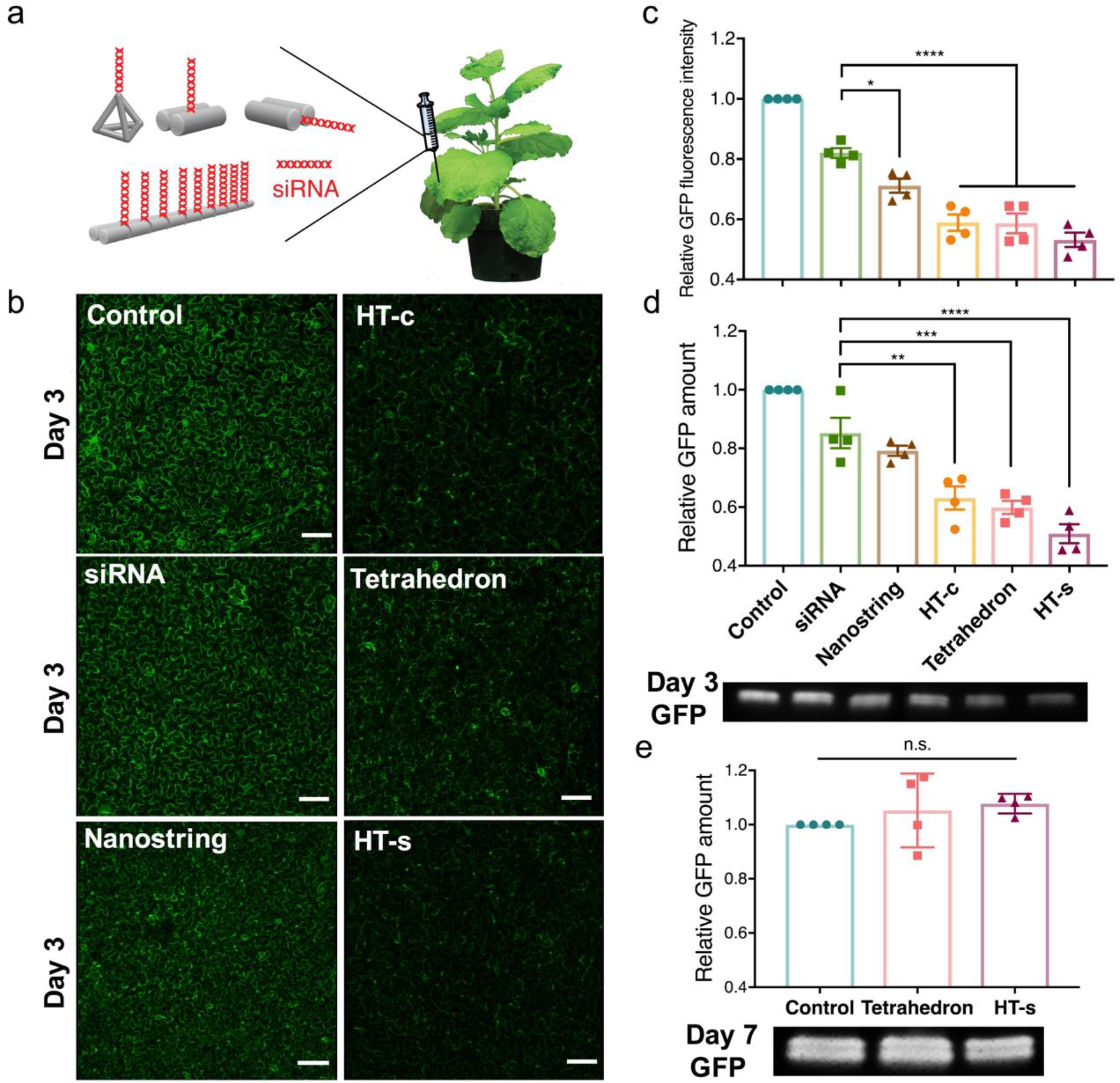
Transient gene silencing with siRNA tethered on DNA nanostructures. GFP silencing efficiency of siRNA-linked nanostructures quantified by confocal imaging and western blotting. a) Infiltration of siRNA linked DNA nanostructures into mGFP5 *Nicotiana benthamiana* leaves. b) Representative confocal images of leaves infiltrated with siRNA-nanostructures 3-days post-infiltration, with non-treated control leaves. Scale bars, 100 µm. c) Quantitative fluorescence intensity analysis of confocal images. *P = 0.0151 and ****P < 0.0001 in one-way ANOVA. d) Representative western blot gel and statistical analysis of GFP extracted from nanostructure treated leaves 2-days post-infiltration. ** P = 0.0013, *** P = 0.0003 and **** P < 0.0001 in one-way ANOVA. e) Representative western blot gel of GFP extracted from leaves treated with siRNA linked to tetrahedron or HT-s nanostructures 7-days post-infiltration. Control vs. tetrahedron = not significant (p = 0.5806), Control vs. HT-s = not significant (p = 0.3444). All error bars indicate s.e.m. (n = 4).

### siRNA attachment locus on nanostructures affects endogenous gene silencing pathways

siRNA mediated gene silencing in plants is a well-known sequence-specific gene regulation mechanism. However, RNA silencing can undergo different gene silencing pathways: transcriptional gene silencing (TGS) or post-transcriptional gene silencing (PTGS). Specifically, PTGS employs microRNA and siRNA pathways for mRNA cleavage or translation repression, while TGS instead undergoes a RNA-directed DNA methylation pathway to inhibit protein translation (48). We tested the siRNA silencing mechanism of DNA nanostructures as illustrated in Figure 4a. Because degradation of transcriptional mRNA is the typical mechanism for gene silencing with exogenously introduced siRNA, we quantified changes in GFP mRNA with quantitative polymerase chain reaction (qPCR). GFP mRNA of mGFP5 *Nicotiana benthamiana* leaf tissues infiltrated with siRNA-linked nanostructures were quantified with qPCR 2-days post-infiltration. Interestingly, as shown in Figure 4b, only siRNA alone and siRNA tethered to the tetrahedron showed a significant (22.3 ± 2.2 % and 50.3 ± 4.9 %, respectively) reduction in GFP mRNA. In contrast, the siRNA tethered to HT monomer or nanostring nanostructures showed a significant *increase* in GFP mRNA of 59.1± 6.5 % for the side-linked monomer HT-s, 45.2 ±1.9 % for the center-linked monomer HT-c, and 35.1± 3.2 % for the nanostring. We further confirmed that the DNA nanostructure alone (HT monomer) doesn’t induce a change in leaf GFP mRNA levels (Figure S14).

The above results suggest that siRNAs conjugated to different DNA nanostructures employ different silencing pathways. Tetrahedron-mediated gene silencing appears to undergo an mRNA-targeted degradation pathway as does free siRNA, while siRNAs linked to either locus on the HT monomer may undergo translation inhibition based on the observed increase and accumulation in mRNA (Figure 4a). Of note, the observed trend of increasing GFP mRNA was consistent with the silencing efficiency trends of the three nanostructures: the side-linked monomer (HT-s) showed the largest mRNA increase and also the largest GPF decrease as measured by western blotting and confocal microscopy. We thus hypothesize that steric and conformational hindrance of the siRNA, determined by the siRNA attachment locus, affects the gene silencing pathway. Specifically, we find that siRNA tethered to the 1D nanostructures (HT monomer, nanostring) have greater steric hindrance than when tethered to the apex of the 0D tetrahedron nanostructure.

To further probe the effect of siRNA linking geometry on gene silencing and to test the above hypothesis, we designed an assay to probe siRNA silencing efficiency of the GFP targeting siRNA under two different linking geometries. Because SWCNT have been previously shown to internalize passively into plant cells (13, 43), we attached siRNA to the surface of a 1D SWCNT but with two different attachment configurations: As shown in Figure 4c, siRNA was either tethered to the surface of a 1D SWCNT (RNA-SWCNTs hybridized), or reversibly loaded on the SWCNT in a releasable manner (RNA-SWCNTs adsorbed). Both constructs were introduced into mGFP5 *Nicotiana benthamiana* leaves and assessed for GFP silencing efficiency (methods). As shown in Figure 4d and Figure S15, both siRNA attachment configurations show similar levels of GFP decrease as quantified by western blotting 2-days post-infiltration: siRNA hybridized to SWCNT decreased GFP expression by 54.3 ± 1.7 % (mean ± SD), and siRNA absorbed onto but releasable from the 1D SWCNT decreased GFP expression by 48 ± 4.8 % (mean ± SD). However, qPCR assessment reveals that GFP mRNA increases by 48.4 ± 8.5 % if the siRNA is hybridized to the SWCNT, whereas the GFP mRNA decreases by 92 ± 1.0 % if the siRNA is releasable from the SWCNT surface (Figure 4e). These results suggest both the silencing efficiency and silencing pathway are affected by the siRNA loading geometry on the nanostructure carrier and the availability of siRNA to the requisite endogenous gene silencing proteins.

**Figure 4.**
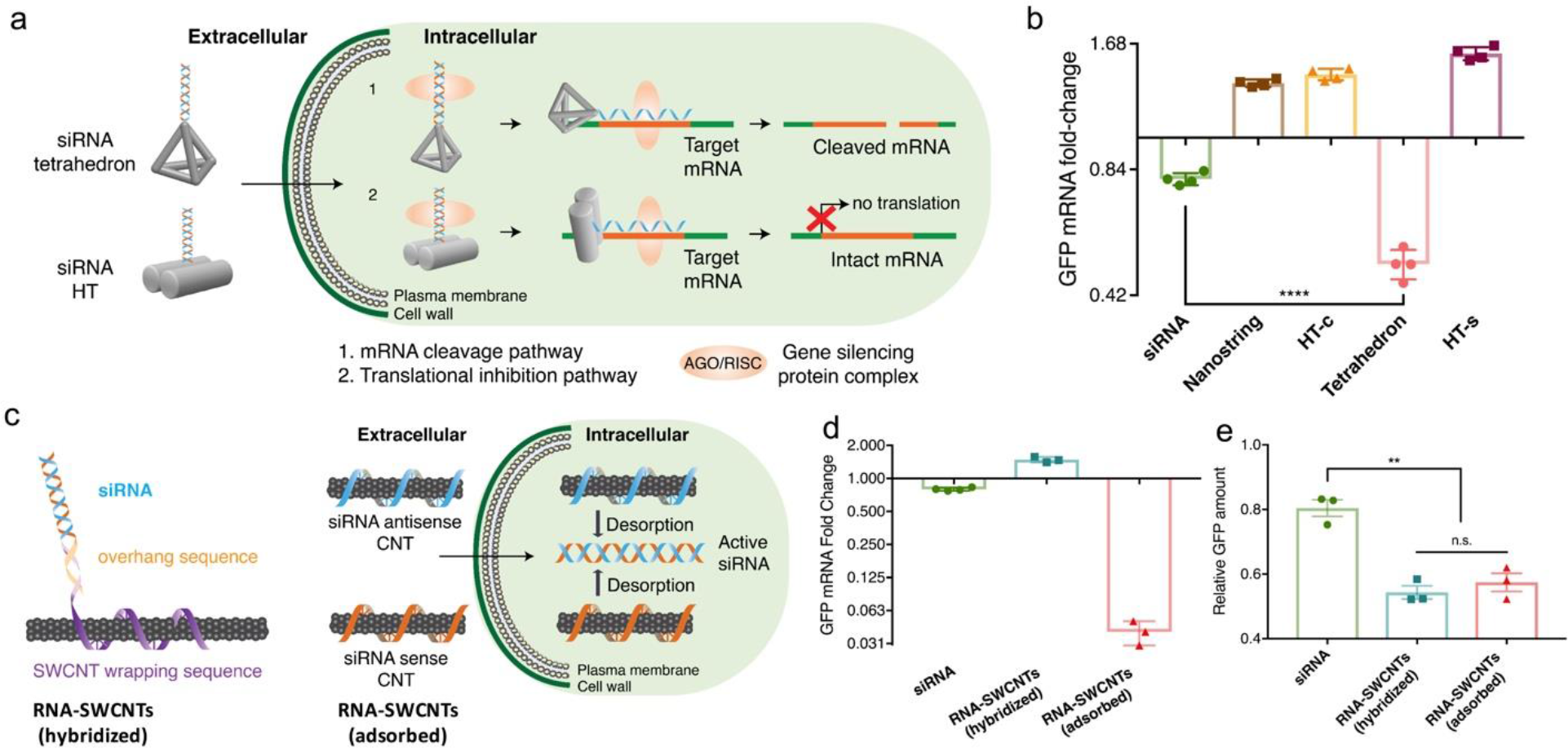
Gene silencing pathways for siRNA-linked nanostructures. a) Proposed silencing pathways induced by siRNA conjugated to DNA nanostructures. b) qPCR of leaves infiltrated with free siRNA, siRNA conjugated nanostring, HT-c, tetrahedron, or HT-s nanostructures 2-days post-infiltration. P **** < 0.0001 in one-way ANOVA. Error bars indicate s.e.m. (n = 4). c) Hybridization or adsorption-based siRNA loading on SWCNTs. d) GFP extracted from siRNA-SWCNT treated leaves 2-days post-infiltration as measured with western blotting. **P = 0.0041, n.s.: non-significant in one-way ANOVA. Error bars indicate s.e.m. (n = 3). e) qPCR of leaves infiltrated with free siRNA, hybridized RNA-SWCNTs, and adsorbed RNA-SWCNTs 2-days post-infiltration, error bars indicate s.e.m. (n = 3).

### DNA nanostructure toxicity analysis

DNA nanostructures have been broadly employed for intracellular delivery in mammalian cells due to their unique programmability, and their inherent biocompatibility compared to inorganic nanomaterials (22). Most prior work comes to a consensus that DNA based nanostructures do not exhibit toxicity towards mammalian systems (24, 49). Since DNA nanostructures have not been used in plant systems to-date, we assessed whether DNA nanostructures are biocompatible in plants. Specifically, we chose to monitor the expression of the respiratory burst oxidase homolog B (NbrbohB) gene, which is a known stress gene upregulated under broad types of stress (mechanical, light, heat, biotic) in *Nicotiana benthamiana* plants (50). Following infiltration of *benthamiana* leaves with either PBS buffer, HT monomer, tetrahedron, or nanostring nanostructures, mRNA of NbrbohB was measured and normalized with respect to the Elongation Factor 1 (EF1) gene, a common *Nicotiana benthamiana* housekeeping gene. As summarized in Figure S16, leaves infiltrated with either of the DNA nanostructures does not result in NbrbohB gene upregulation compared to adjacent areas within the same leaf treated only with PBS buffer. Furthermore, the structural integrity of the plant cells is unperturbed by introduction of the various DNA nanostructures (Figure 2 and 3, Figure S3, S4 and S6). Our analyses suggest that DNA nanostructures do not induce a stress response in plants and are a biocompatible mode of siRNA delivery to plant cells.

## Discussion

DNA nanostructures have been extensively studied in animal systems for cell internalization, intracellular delivery, and for downstream diagnostic and therapeutic applications owing to their unique sequence-structure programmability and inherent biocompatibility (23–25, 51, 52). Prior work has focused on studying DNA nanostructure internalization in mammalian cells, and has revealed that nanostructure size, shape, charge, and stiffness or compactness can influence the cellular internalization and uptake pathway (34, 51, 53, 54). Analogous work in plant systems is lacking, where a few studies have reported the biolistic or passive uptake, translocation, or localization of engineered nanoparticles (carbon nanotubes, SiO_2_, quantum dots, TiO_2_ NPs) to plants (13, 14, 28, 32, 33, 55), while DNA nanostructure use in plants remains unexplored. Orthogonally, gene silencing through the introduction of siRNA has become a broadly-adopted tool to inactivate gene expression, to probe biosynthetic pathways, and to serve as exogenous regulators of developmental and physiological phenotypes in plants (56, 57).

Herein, we demonstrate that DNA nanostructures can effectively be designed to internalize into plant cells through passive infiltration, and that siRNA can be controllably tethered to specific loci on the DNA nanostructures for effective gene silencing in *Nicotiana benthamiana* leaves. We show that siRNA delivered by DNA nanostructures silences a transgene more effectively than siRNA delivered alone. We further find that the structural and mechanical properties (size, shape, compactness, and stiffness) of DNA nanostructures, and siRNA conjugation loci, affect not only nanostructure internalization into plant cells, but also subsequent gene silencing efficiency, and gene silencing pathway preference for either TGS or PTGS. Once intracellular, siRNAs are processed by nucleases for assembly into the RNA-induced silencing complex (RISC) (58). The RISC complexes with a single-strand of the siRNA duplex and guides the RISC complex to its complementary mRNA target where it will cleave, and thus inactivate, the target mRNA, effectively silencing the downstream protein product. Several forms of RISC that differ in size and composition have been reported (59, 60), and are presumed to undergo mechanistic variations in their silencing pathways. While TGS accomplishes gene silencing via RNA-directed DNA methylation, PTGS accomplishes gene silencing through siRNA-guided mRNA cleavage or translational inhibition(48).

In this work, we find that the likely gene silencing mechanisms undertaken by siRNA linked to DNA nanostructures depend on the siRNA attachment locus and steric availability of the attached siRNA. Interestingly, siRNA tethered to 0D nanostructures show gene silencing both at the transcript (mRNA) and protein levels, whereby siRNA attached to 1D nanostructures shows gene silencing at the protein level, but shows an increase in mRNA transcript levels. This phenomenon of increased mRNA implies a possible silencing pathway and mechanism for siRNA delivered with HT or nanostring DNA nanostructures, in which translational inhibition of GFP expression is preferred over direct mRNA cleavage. We hypothesize that protein translation inhibition leads to continuous production and accumulation of repressed mRNAs, as we observe through qPCR of leaves treated with select nanostructure carriers. Specifically, we hypothesize that the steric accessibility of siRNA conjugated to different DNA nanostructures by endogenous silencing proteins plays a dominant role in determining the silencing mechanism, whereby formation of the RISC protein complex that leads to mRNA cleavage may be hindered by the proximity of a nanostructure scaffold for 1D nanostructures, but absent for small 0D nanostructures.

In summary, DNA nanostructures can serve as effective scaffolds and nanoscale vehicles for siRNA delivery to plants for efficient gene silencing. This work establishes DNA nanostructures as a programmable toolset for the delivery of exogenous biomolecules such as siRNA to plants, and establishes guidelines for the design of DNA nanostructures for effective uptake into plant cells for various applications in plant biotechnology.

## Acknowledgements

We acknowledge support of the Burroughs Wellcome Fund (CASI), a USDA AFRI grant with Award # 2018-67021-27964, an FFAR New Innovator Award, and an NSF-USDA-BBSRC grant. H.Z. acknowledges the support of the Chinese National Natural Science Foundation (Grant 21605153), G.S.D. is supported by the Schlumberger Foundation. We acknowledge the Berkeley Molecular Imaging Center, and the QB3 Shared Stem Cell Facility.

## Author contributions

H.Z. and M.P.L. conceived the project, designed the study, and wrote the manuscript. H.Z. conducted the experiments. H.Z. and G.S.D. analyzed the data. H.L.Z. performed AFM imaging, mechanical simulation and calculation, and analyzed the data. G.S.D and T.Y. performed the q-PCR, Western Blot and confocal imaging experiments. N.S.G, F.J.C, A.J.A., and C.F. discussed the project and edited the manuscript. All authors have edited and commented on the manuscript, and have given their approval of the final version.

## Competing interests

The authors declare no competing interests.

Correspondence and requests for materials should be addressed to M.P.L.

## Materials and Methods

### Plant growth

Transgenic mGFP5 *Nicotiana benthamiana* seeds (obtained from the Staskawicz Lab, UC Berkeley) were germinated and kept in SunGro Sunshine LC1 Grower soil mixture for four weeks before use. Plants were allowed to mature to 3-4 weeks of age within the chamber before experimental use.

### Infiltration of leaves with nanomaterials (SWCNTs and DNA nanostructures)

4 week old mGFP5 *Nicotiana benthamiana* plants were punctured on the abaxial surface of the leaf lamina with a razor, and 100 μL of DNA nanostructure solutions were infiltrated through the puncture with a 1 mL needleless syringe. Post-infiltration, leaves were left in plant pots and analyzed after above-mentioned periods of time (2-3-days or 7-days post-infiltration) to quantify nanostructure internalization or GFP gene silencing efficiency. Nanostructure internalization efficiencies (calculated with colocalization analysis), and gene silencing efficiency (calculated with quantitative GFP fluorescence intensity analysis, qPCR, and western blotting) were subsequently performed as described in supplementary methods.

### Toxicity analysis of nanostructures in plants

The expression of an oxidative stress gene (NbRbohB) in *Nicotiana benthamiana* leaves was measured to test for plant stress and toxicity through qPCR (with primers fNbRbonB and rNbRbonB, sequences in Table S1). EF1 was measured as a housekeeping gene. An annealing temperature of 60°C was used for qPCR, which was run for 40 cycles, and the ddCt method was used to obtain the normalized NbRbohB expression-fold change with respect to the EF1 housekeeping gene and control sample.

