## Supplemental Information for "DNA Nanostructures Coordinate Gene Silencing in Mature Plants"

[1] Department of Chemical and Biomolecular Engineering, University of California, Berkeley, CA 94720, USA.
[2] Department of Molecular and Cell Biology, University of California, Berkeley, CA 94720, USA.
[3] Division of Physical Biology & Bioimaging Center, Shanghai Synchrotron Radiation Facility, CAS Key Laboratory of Interfacial Physics and Technology, Shanghai Institute of Applied Physics, Chinese Academy of Sciences, Shanghai 201800, China.
[4] School of Chemistry and Chemical Engineering, Shanghai Jiao Tong University (SJTU), Shanghai, 200240, China.
[5] Innovative Genomics Institute (IGI), Berkeley, CA 94720, USA.
[6] California Institute for Quantitative Biosciences, QB3, University of California, Berkeley, CA 94720, USA.
[7] Chan-Zuckerberg Biohub, San Francisco, CA 94158, USA.
[§] Authors contributed equally to this work.

[*]

**Supplementary Table S1** | Sequences of oligonucleotides used in this study.

| Name | Sequence (5'-3') |
|---|---|
| **H1** | GGACTTGTAGCGATACGACTCCGACGAGACTAGTAACTCTTG |
| **H2** | GGATGCGGAATGACAGCTACAAGTCCCAAGAGTTACGCTCTCCATTC |
| **H3** | TGTCACAGTAAGTCTTGTCATTCC |
| **H4** | GGGCTTGAATGGAGAGCCATCACTCATGTGAACCCATGAGTGATGTAGTCTCGTCGGAGTCGTATCAGACTTACT |
| **H5** | ACGAGACTACATGGTCAGATTCGTAGGTCCGATACGACTCCG |
| **H6** | GCATCCGATCCGTCCTGTCGGACCTACGAATCTGACCACCGAGAATC |
| **H7** | AAGCCCGATTCTCGGTCACTCATGGGTTCA |
| **H8** | CATGAGTGATGTAGTCTCGTCGGAGTCGTATCAGACTTACTGTGACAAGTAAGTCTGACAGGACGGATC |
| **I** | CATGAGTGATGTAGTCTCGTCGGAGTCGTATCAGACTTACTGTGACA |
| **Bio-H1** | Biotin-GGACTTGTAGCGATACGACTCCGACGAGACTAGTAACTCTTG |
| **Cy3-H1** | Cy3-GGACTTGTAGCGATACGACTCCGACGAGACTAGTAACTCTTG |
| **H1-15-RNA** | TAC ACG CAT CCT TAG GGACTTGTAGCGATACGACTCCGACGAGACTAGTAACTCTTG |
| **H2-15-RNA** | TAC ACG CAT CCT GGA TGC GGA ATG ACA TGC TAC AAG TCC CAA GAG TTA CGC TCT CCA TTC |
| **H5-15 -CNT** | ACGAGACTACATGGTCAGATTCGTAGGTCCGATACGACTCCG TAC ACG CAT CCT TAG |
| **A** | GAGCGTT A GCCACAC A CACAGTC |
| **B** | TTAGGCG A GTGTGGC A GAGGTGT |
| **C** | CGCCTAA A CAAGTGG A GACTGTG |
| **D** | AACGCTC A CCACTTG A ACACCTC |
| **A-15-RNA** | G CAT CCT TAG AAA AAA GAGCGTT A GCCACAC A CACAGTC |
| **sense-15** | CTA AGG ATG CGT GTA GGU GAU GCA ACA UAC GGAA TT |
| **antisense** | UUC CGU AUG UUG CAU CACC |
| **$(GT)_{15}$-15** | GTGTGTGTGTGTGTGTGTGTGTGTGTGTGT CTA AGG ATG CGT GTA |
| **$(GT)_{15}$-15-Cy3** | GTGTGTGTGTGTGTGTGTGTGTGTGTGTGT CTA AGG ATG CGT GTA-Cy3 |
| **fGFP** | AGTGGAGAGGGTGAAGGTGATG |
| **rGFP** | GCATTGAACACCATAAGAGAAAGTAGTG |
| **fEF1** | GCATTGAACACCATAAGAGAAAGTAGTG |
| **rEF1** | ACGCTTGAGATCCTTAACCGCAACATTCTT |
| **fNbrbohB** | TTTCTCTGAGGTTTGCCAGCCACCACCTAA |
| **rNbrbohB** | GCCTTCATGTTGTTGACAATGTCTTTAACA |

**Supplementary Table S2** | Calculation of nanostructure parameters

<table>
<tr><th>Structures</th><th>Width (nm)</th><th>Length (nm)</th><th>Relative stiffness ($k_b$structure/$k_b$ds $_{DNA}$)</th><th>Compactness ($C_{sphere}$/$C_{structure}$)</th><th colspan="2">Aspect ratio after conjugated with siRNA</th></tr>
<tr><td>Tetrahedron</td><td>2.4</td><td>2.4</td><td>×</td><td>0.55</td><td colspan="2">5:1</td></tr>
<tr><td rowspan="2">Hairpin –tile (HT)</td><td rowspan="2">5</td><td rowspan="2">16</td><td rowspan="2">0.83</td><td rowspan="2">0.45</td><td>Center</td><td>1:1</td></tr>
<tr><td>Side</td><td>5:1</td></tr>
<tr><td>Nanostring</td><td>5</td><td>320</td><td>$1.0 \times 10^{-4}$</td><td>0.11</td><td colspan="2">20:1</td></tr>
<tr><td>SWCNTs</td><td>1</td><td>~500</td><td>GPa to TPa(1, 2)</td><td>×</td><td colspan="2">35:1</td></tr>
</table>

The relative bending stiffness of a 1D structure was calculated based on a beam-shaped model and the compactness of a 3D structure was defined by the ratio area$^{1.5}$/volume, which is dimensionless and minimized by a sphere (see details in Methods).

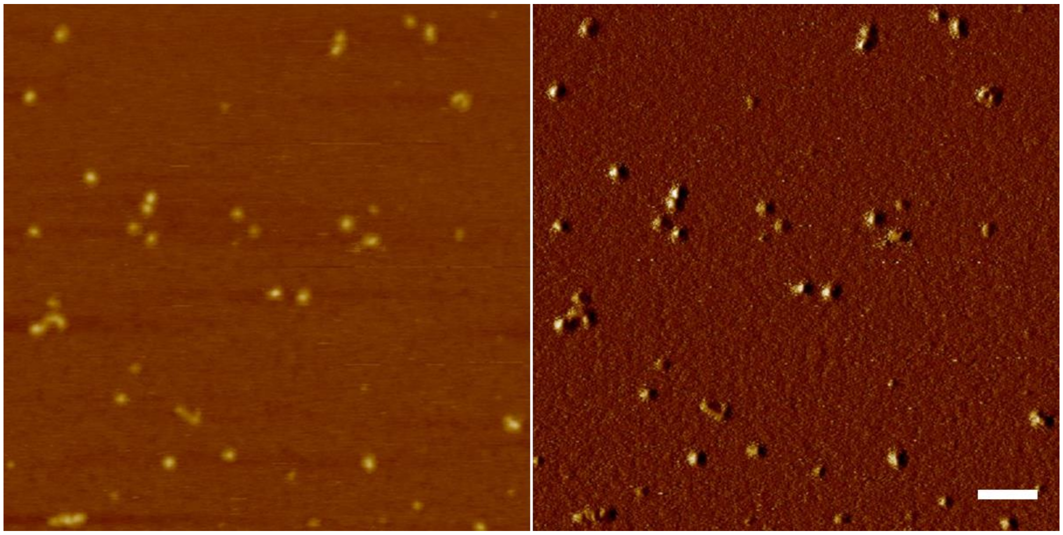

**Figure S1.** AFM images of tetrahedron nanostructures. Left: height image; Right: phase image. Scale bar: 100 nm.

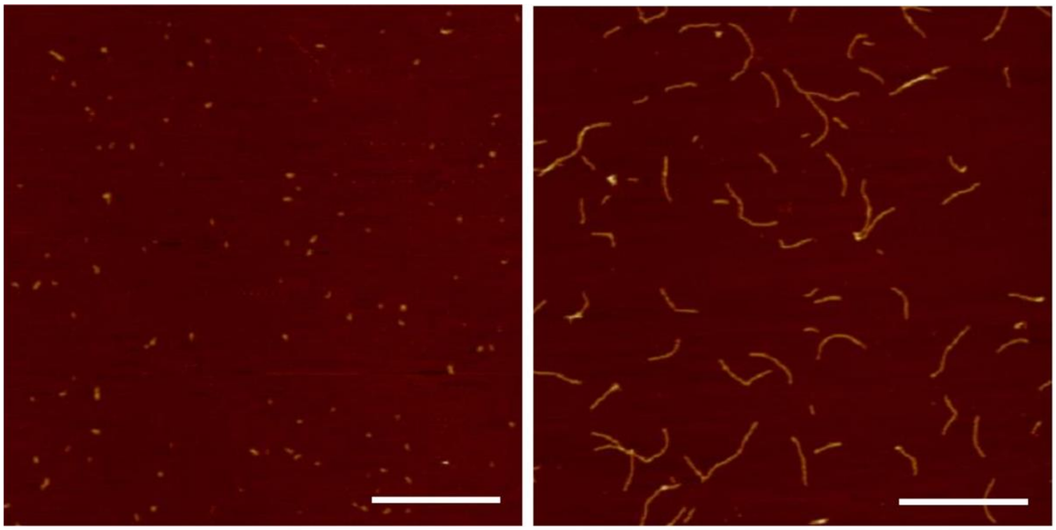

**Figure S2.** AFM height images of HT monomer (left) and nanostring (right) nanostructures. Scale bar: 500 nm.

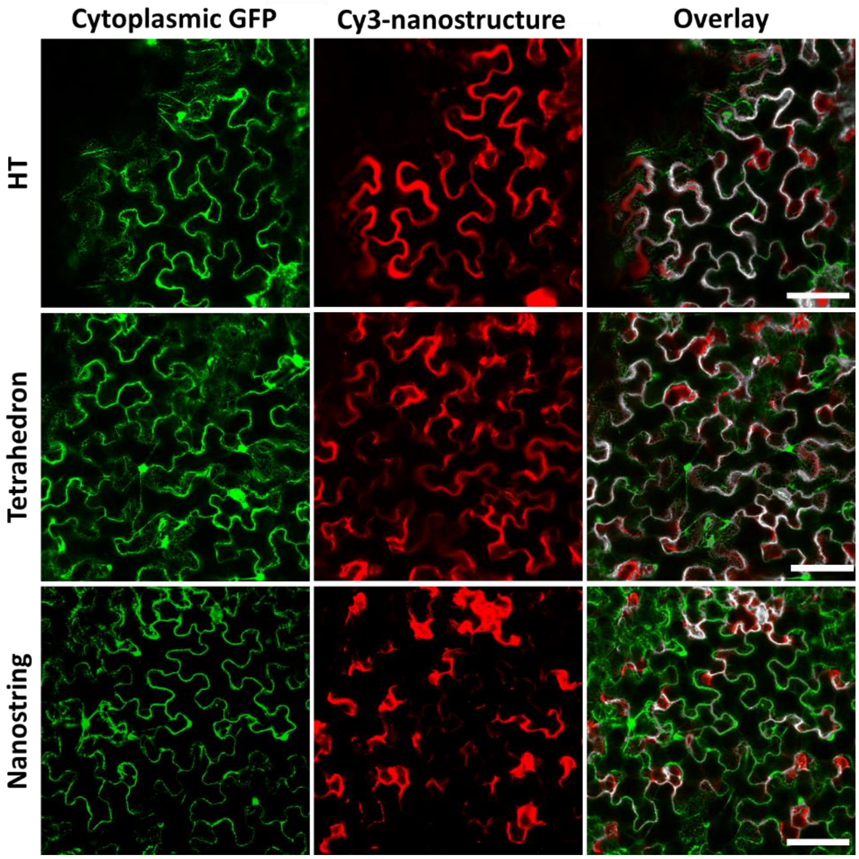


**Figure S3.** Representative confocal images of Cy3 labeled HT monomer, tetrahedron, and nanostring colocalized with the GFP cytoplasm of mGFP5 *benthamiana*. Scale bars, 100 µm.

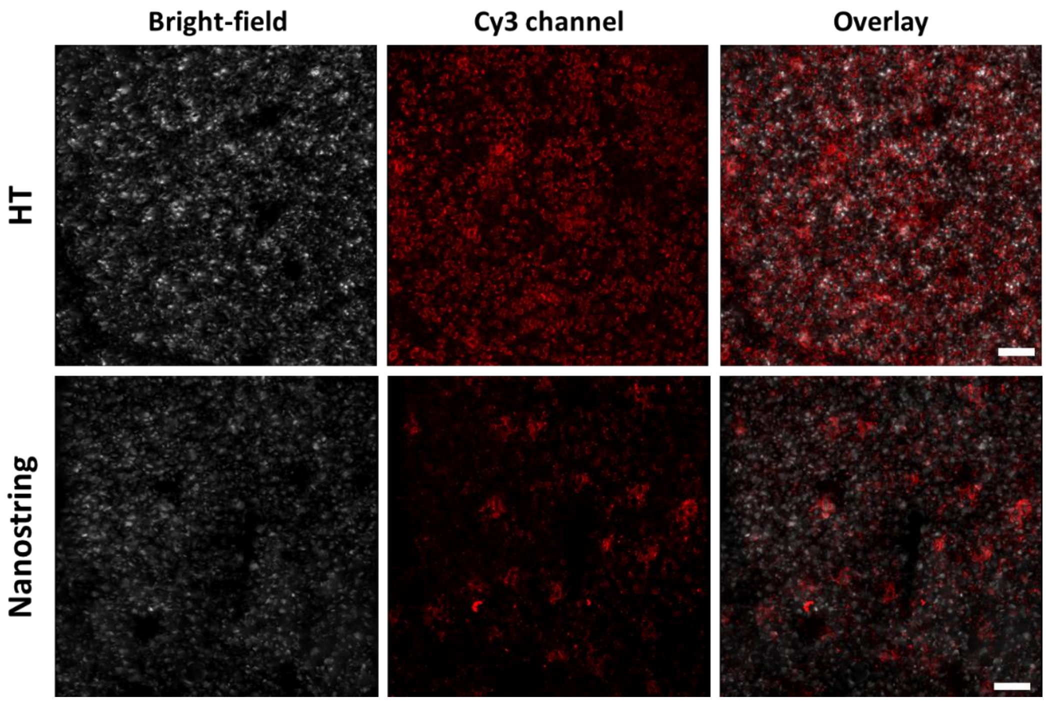


**Figure S4.** Representative confocal images of Cy3 labeled HT monomer, and nanostring. Scale bars, 100 µm.

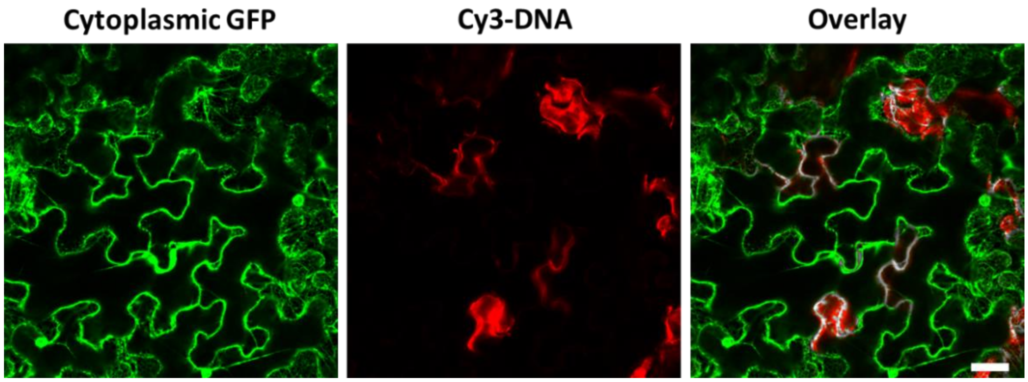


**Figure S5.** Representative confocal images of Cy3 labeled single stranded DNA colocalized with the GFP cytoplasm of mGFP5 *benthamiana*. Scale bars, 50 µm.

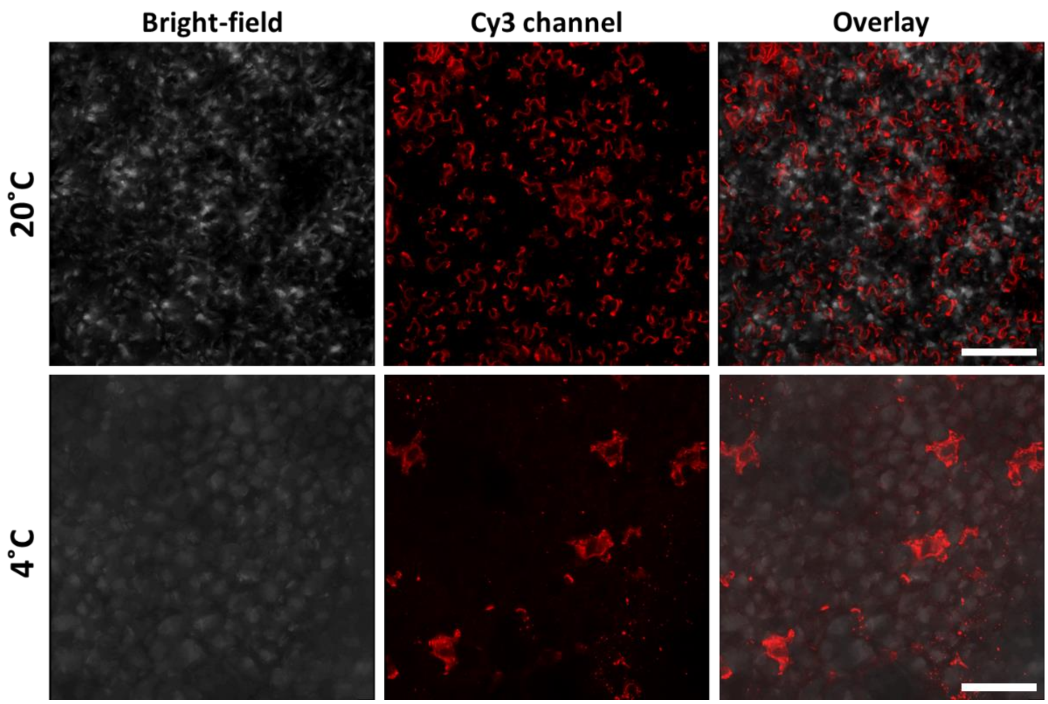


**Figure S6.** Representative confocal images showing the temperature dependence of nanostructure internalization for the Cy3 labeled HT monomer. Scale bars, 50 µm.

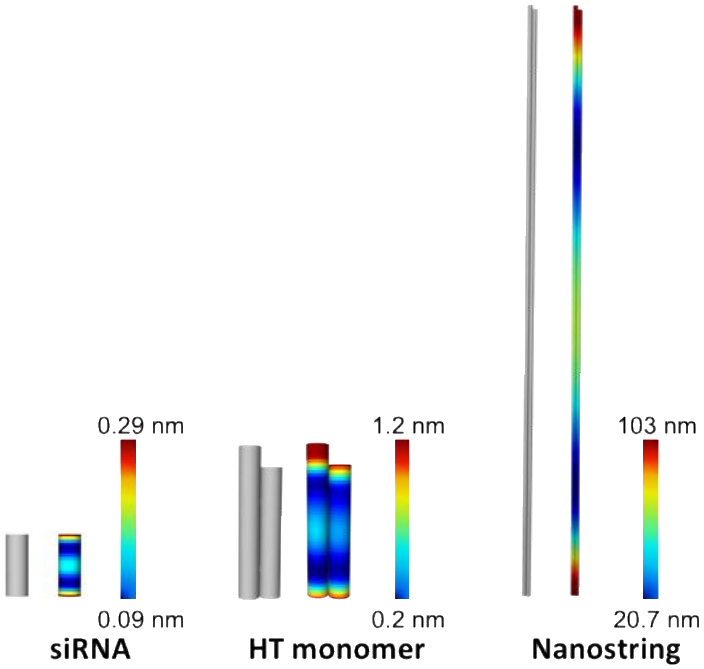


**Figure S7.** Equilibrium conformation and heat map color range of root-mean-square fluctuations (RMSF) for the siRNA, HT monomer, and nanostring, simulated by CanDo (3, 4). Blue and red represent low and high relative flexibility, respectively. Bluest = 0% RMSF and reddest = 95% RMSF.

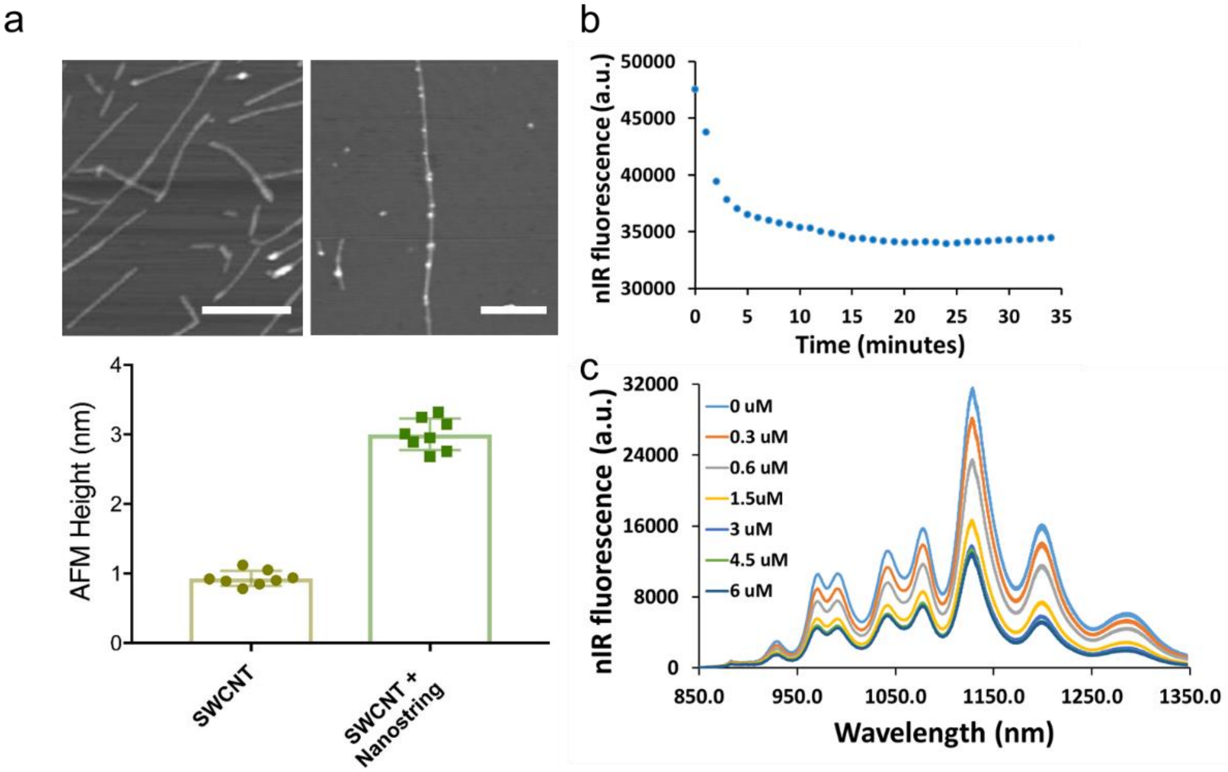


**Figure S8.** a) Representative AFM images of single-walled carbon nanotubes (SWCNT, upper left), SA-biotin nanostring conjugated SWCNT (upper right), and statistical height analysis of SWCNT (~1 nm) and SWNT-nanostring conjugation (~3 nm). Scale bars, 100 nm. b) SWCNT nIR fluorescence change with time when nanostrings hybridize to SWCNTs. c) SWCNT nIR spectra after adding different nanostring concentrations, each spectrum was taken 10 minutes after nanostring addition.

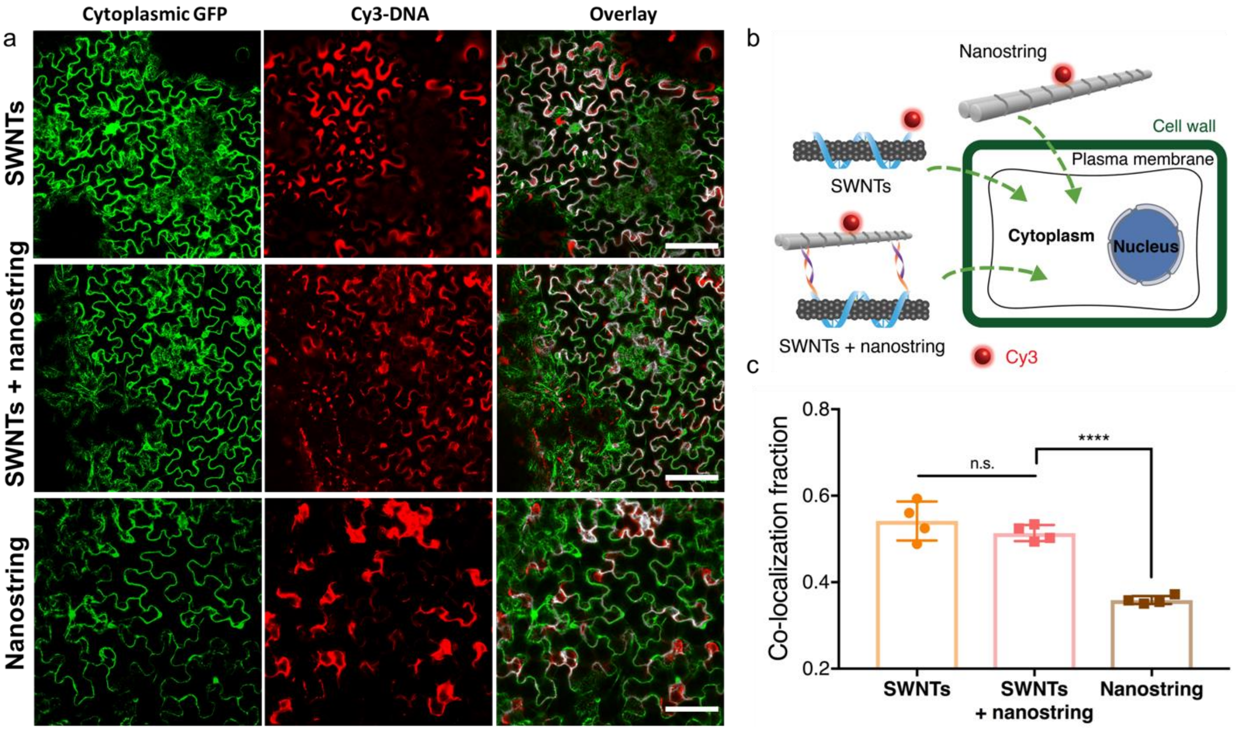


**Figure S9.** a) Representative confocal images showing different internalization behaviors of SWCNTs (top), nanostring conjugated SWCNTs (middle), and nanostring nanostructures alone (bottom) post-infiltration in mGFP5 *benthamiana*. Scale bars, 100 µm. b) Schematic depicting proposed the internalization behaviors of three different nanostructures post-infiltration into plant cells: SWCNTs wrapped by Cy3 labeled $GT_{15}$ ssDNA, nanostrings labeled with Cy3 at the nanostring center and hybridized onto SWCNT, and the Cy3 labeled nanostring alone. c), Colocalization analysis of Cy3 fluorescence (indicating nanostructure) with the GFP fluorescence (indicating plant cell cytosol) after 12 (nanostring) or 6 (SWCNTs and SWCNTs + nanostring) hours post-infiltration. P**** < 0.0001 in one-way ANOVA, n.s.: not significant difference. Error bars indicate s.e.m. (n = 4)

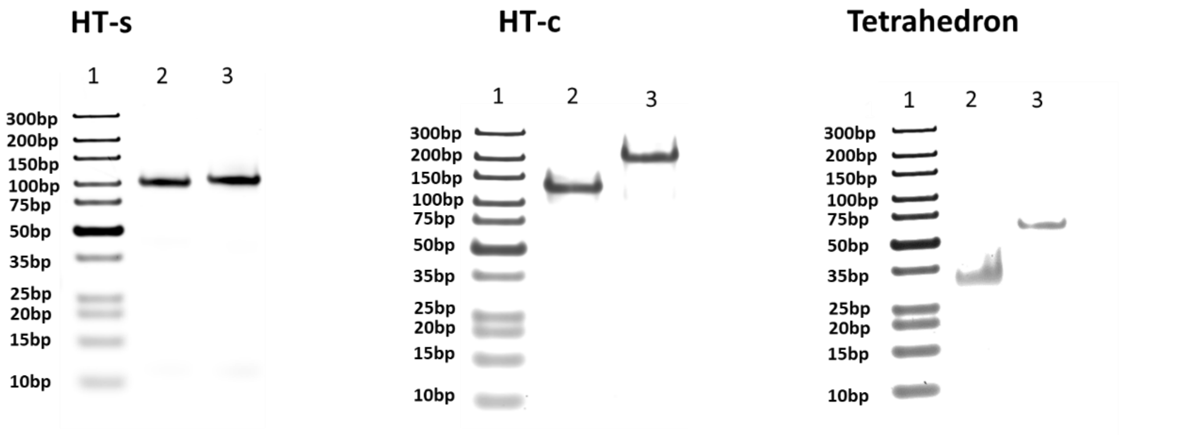


Lane 1: Marker, 3μL
Lane 2: HT-s, 1μM, 5μL
Lane 3: HT-s + siRNA, 1μM, 5μL

Lane 1: Marker, 3μL
Lane 2: HT-c, 1μM, 5μL
Lane 3: HT-c+ siRNA, 1μM, 5μL

Lane 1: Marker, 3μL
Lane 2: tetrahedron, 1μM, 3μL
Lane 3: lane 2 tetrahedron + 1 siRNA, 1μM, 3μL

**Figure S10.** 10% Native-PAGE gels to verify attachment of siRNA to HT monomer and tetrahedron at different loci.

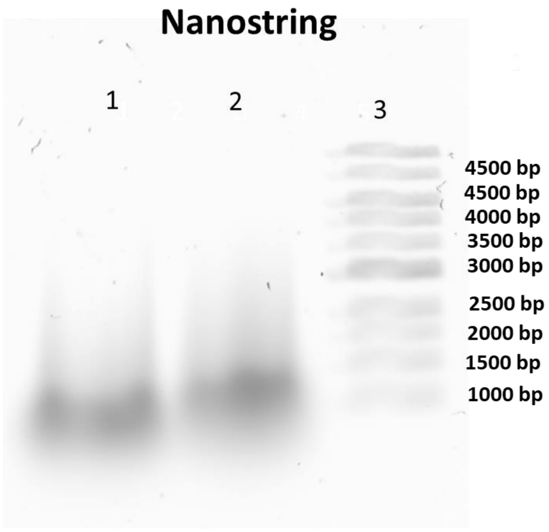


Lane 1: Nanostring 1 μM, 2 μL
Lane 2: Nanostring + siRNA, 1 μM, 2 μL
Lane 3: Marker, 2 μL

**Figure S11.** 1% agarose gel confirming attachment of siRNA to nanostring.

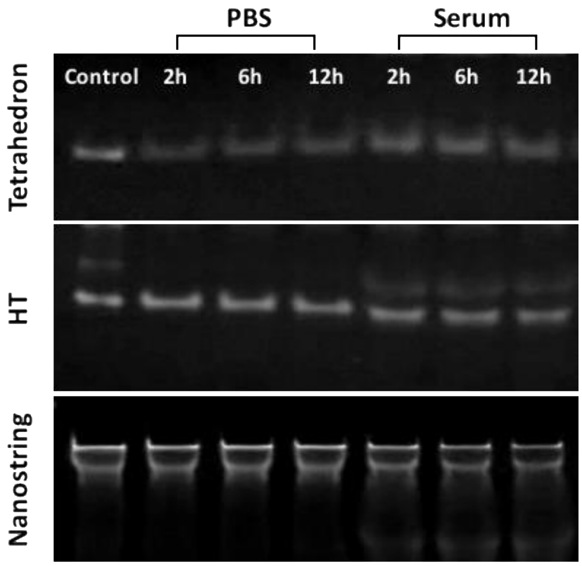


**Figure S12.** 10% Native PAGE gels showing the stability of the DNA nanostructures in different media as a function of incubation time.

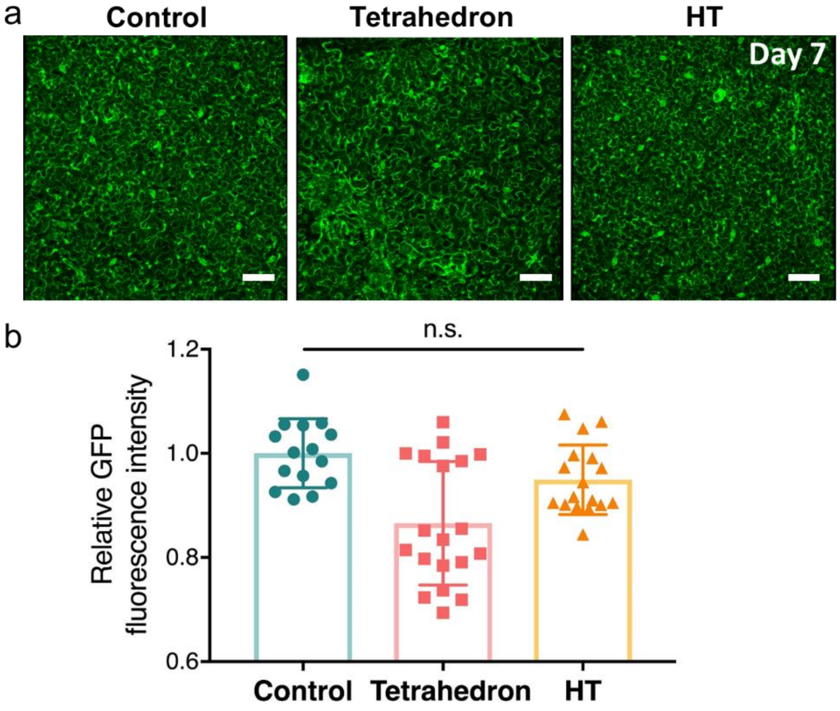


**Figure S13.** a) Representative confocal images of mGFP5 *benthamiana* leaves 7-days post-infiltration with PBS, siRNA-tetrahedron nanostructures, or siRNA-HT momomer nanostructures, showing GFP fluorescence recovery. Scale bars, 100 µm. b) Quantitative fluorescence intensity analysis of confocal images. n.s.=non-significant (s.d. n = 15).

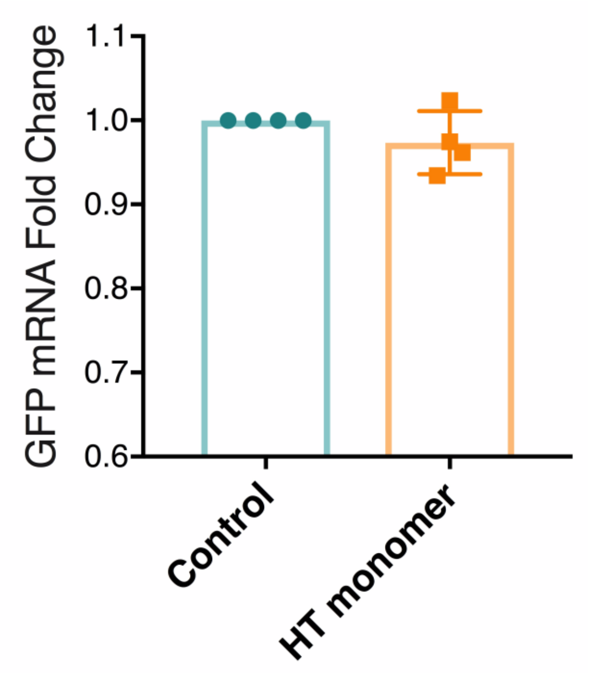


**Figure S14.** qPCR results for leaves infiltrated with HT monomer alone after 2-days post-infiltration, showing no mRNA change. Error bars indicate s.e.m. (n = 4).

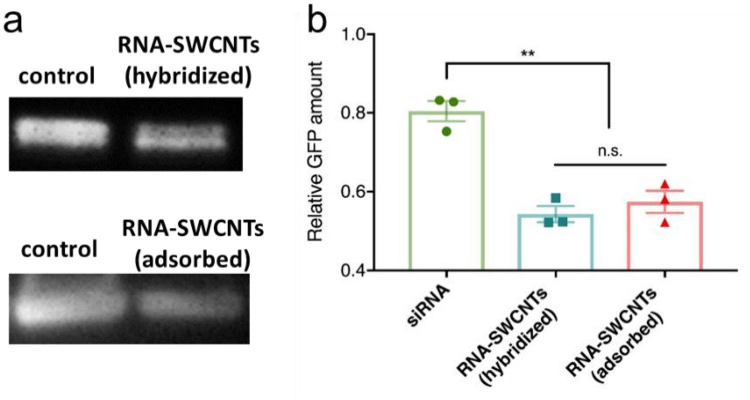


**Figure S15.** a) Representative western blot gel and analysis showing GFP extracted from free siRNA, siRNA attached SWCNTs through hybridization, or siRNA adsorbed to SWCNT 2-days post-infiltration to mGFP5 *benthamiana*. b) Statistical analysis of the results showing GFP proteins extracted from different SWCNT-treated leaves 2-days post-infiltration. P** = 0.0041 in one-way ANOVA. Error bars indicate s.e.m. (n = 3).

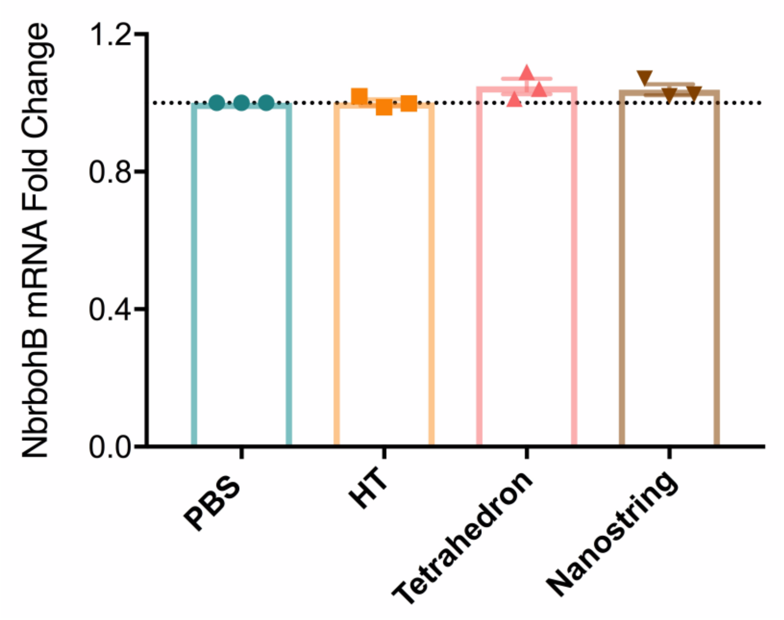


**Figure S16.** Quantitative qPCR analysis of NbrbohB, a known stress gene, to test the toxicity of the DNA nanostructures used to deliver siRNA. Error bars indicate s.e.m. (n = 3).

**Supplementary Video S1.** Mechanical stiffness calculations quantify the root mean square fluctuations (RMSF) of HT the monomer nanostructure simulated by CanDo (3, 4). White and red represent low and high relative flexibility, respectively.

**Supplementary Video S2.** Mechanical stiffness calculations quantify the root mean square fluctuations (RMSF) of HT the nanostring nanostructure simulated by CanDo (3, 4). White and red represent low and high relative flexibility, respectively.

**Calculation of relative nanostructure bending stiffness**

The bending stiffness ($k_b$) of a beam-shaped structure is described by:

$$k_b = \frac{3EI}{L^3}$$

where $E$ is the Young's modulus (elastic modulus),

$L$ is the length of the DNA nanostructure,

$I$ is the area moment of inertia.

To estimate the moment of inertia, we treat a 1D nanostructure as a bundle of *N* rigidly linked cylindrical rods of radius *r*, where *I* can be calculated in terms of *I (61, 62)*. The moment of inertia of each dsDNA helix with respect to its own center of mass is *i*, and is displaced from the nanostructure's center of mass by a distance *R*. By the parallel axis theorem, the moment of inertia of the dsDNA helix with respect to the nanostructure's center of mass is $i+MR^2$, where *M* is the mass of the helix. Assuming uniform density,

$$i = \frac{1}{2}Mr^2$$

and thus

$$I = N(i + MR^2) = N\left(i + 2i\frac{R^2}{r^2}\right)$$

with Relative stiffness (see Table S2)

$$\frac{k_b(nanostructure)}{k_b(dsDNA)} = \frac{I}{i} \times \left(\frac{L(dsDNA)}{L(nanostructure)}\right)^3 = N\left(1 + 2\frac{R^2}{r^2}\right) \times \left(\frac{L(dsDNA)}{L(nanostructure)}\right)^3$$

**Structural conformation and mechanical stiffness simulation**

The structural shape and mechanical compliance of siRNA, HT monomer, and nanostring nanostructures were modeled using the finite element model with CanDo (cando-dna-origami.org), to predict the structural shape and mechanical flexibility of the DNA nanostructures. The nanostructures were modeled as homogeneous elastic rods with isotropic bending stiffness. The structural and mechanical parameters of two-node beam finite elements composing each rod are as listed below:

- B-form DNA helix is modeled as a worm-like chain.
- Axial length per base pair: 0.34 nm;
- Helical diameter: 2.25 nm;
- Base pairs per turn: 10.5;
- Bending stiffness: 230 pN·nm$^2$;
- Stretching modulus: 1100 pN;
- Torsional stiffness: 460 pN·nm$^2$.

The backbone bending and torsional stiffness are reduced by 100-fold when there are nicks in the DNA double helix, and thus single-stranded DNA present as sticky ends or loops are modeled as entropic springs using a modified freely jointed chain model. Interhelical crossovers are treated as rigid component with zero length.

The structural conformation and mechanical stiffness of DNA nanostructures at ground-state solution are performed by normal mode analysis (63, 64) with CanDo. The thermal fluctuations are quantified using computing root-mean-square fluctuations (RMSF) of base pairs computed at a temperature of 298 K. Here, RMSF measures the magnitude of motion of base pairs and exhibit flexibility of DNA nanostructures.

**Calculation of nanostructure compactness**

The compactness of a 3D structure relates the enclosing surface area (A) with the volume (V) and can be defined by the ratio area$^{1.5}$/volume, which is dimensionless and can be minimized by a sphere.

The compactness of 3D structure

$$C_{structure} = A^{1.5}/V$$

For a sphere:

$$A_{sphere} = 4\pi r^2$$

$$V_{sphere} = \frac{4}{3}\pi r^3$$

The compactness of sphere

$$C_{sphere} = {A_{sphere}}^{1.5}/V_{sphere}$$

$$=3\sqrt{4\pi}$$

Which is the minimum compactness of a solid since the sphere encloses maximum volume for a constant surface area.

The regular compactness is defined as

$$C = C_{sphere}/C_{structure}$$

$C_{sphere}$ is the calculated value of a sphere, and $C_{structure}$ is the calculated value of the sphere DNA nanostructures.

### Supplementary Materials and Methods

#### Chemicals and materials.

Super purified HiPCO SWCNTs (Lot # HS28-037) were purchased from NanoIntegris, and used for all SWCNT-based experiments. The following chemicals were purchased from Sigma-Aldrich: sodium chloride, potassium chloride, magnesium chloride hexahydrate, bovine serum albumin (heat shock fraction). Single stranded RNA and DNA oligonucleotides were purchased from and purified by Integrated DNA Technologies, Inc. (IDT); DNA oligonucleotides labeled with biotin or Cy3 were purified by HPLC, and dissolved in Milli-Q water before use. The concentration of each strand was estimated by measuring the UV absorbance at 260 nm using a UV-3600 Plus UV-Vis-NIR Spectrophotometer (Shimadzu Scientific Instruments, Columbia, U.S.A.). Streptavidin was purchased from Sigma-Aldrich Co. LLC. UltraPure DNase/RNase-free distilled water from Invitrogen was used for qPCR experiments, and EMD Millipore Milli-Q water was used for all other experiments.

#### Non-denaturing polyacrylamide gel electrophoresis (PAGE).

1 µM, of a 5 µL volume of each assembled sample was loaded onto a 10% PAGE (19:1 acrylamide/bisacrylamide in 1×TAE/$Mg^{2+}$ buffer). Gels were run at 100 V (constant voltage) for two hours with an electrophoresis apparatus (Bio-rad, United States). Next, gels were stained with 1× SYBR® Gold nucleic acid dye (Thermo Fisher Scientific, United States) and scanned with a Typhoon FLA 9500 instrument (GE Healthcare life Sciences, United States of America).

#### AFM characterization.

2-3 µL of the nanostring sample was deposited on a freshly cleaved mica surface and left to adsorb on the surface for 3 minutes. For AFM imaging, the mica surface was slowly rinsed with water three times (each time with 10 µL water) to remove salt. Next, the mica surface was dried with a mild air stream by an ear-washing bulb and was imaged with a MultiMode 8 AFM with NanoScope V Controller (Bruker, Inc.) under tapping mode in air. All AFM images were analyzed by NanoScope Analysis v1.50.

#### Biotin-streptavidin binding assays.

The strand H1-biotin was purchased and synthesized by IDT with biotinylation at the 5' end. After the biotin labeled TH monomer or nanostrings were constructed as described before, a stoichiometric amount of streptavidin in 1×TAE/$Mg^{2+}$ buffer was added, and the final molar ratio of streptavidin to biotin-nanofilament was 10:1. The mixture was left at room temperature for 5 min and then characterized by AFM as described above.

**Self-assembly of DNA tetrahedral nanostructure.**

The oligonucleotides of A, B, C, D (see Table S1 for detailed sequences) were stoichiometrically mixed in 1×TM buffer (20 mM Tris, 50 mM $MgCl_2$, pH 8.0), then assembled through annealing with a PCR machine (Veriti™ 96-Well Thermal Cycler, Thermo Fisher, United States) at 95℃ for 10 min, followed by cooling down to 4℃ in 30 s. The assembled structure was characterized with atomic force microscopy (AFM). For the tetrahedron with 15 nucleotide (nt) overhangs for siRNA conjugation, the A-15 stand (Table S1) was used instead of the A strand.

**Self-assembly of hairpin-tile (HT) monomer and 1D nanostring nanostructures.**

The oligonucleotides of H1, H2, H3, and H4 (see Table S1 for detailed sequences) were stoichiometrically mixed in a 1×TAE/$Mg^{2+}$ buffer containing 40 mM Tris base, 20 mM acetic acid, 2 mM EDTA, and 12.5 mM magnesium acetate (pH 8.0). Next, the DNA strand solution was slowly cooled down from 95˚C to 20℃ over 24 hours in a water bath insulated in a Styrofoam box. To co-polymerize the 1D nanostrings, 1 µM of TH monomers A (H1, H2, H3, and H4) and B (H5, H6, H7, and H8) at equimolar concentrations were mixed with initiator strand I in a 1:0.1 ratio. The mixtures were further incubated at 20℃ for 1 h and were then characterized by AFM. For the HT monomer and nanostring with 15-nt overhangs with siRNA conjugation at the center, the H1-15-RNA stand was used instead of the H1 strand. For HT monomer with 15-nt overhangs with siRNA conjugation at the side, H2-15-RNA stand was used instead of the H2 strand. For nanostring with 15-nt overhangs for SWCNTs conjugation, the H5-15-CNT stand was used instead of the H5 strand.

**Hybridization of DNA nanostructures with double stranded siRNA.**

The duplex siRNA with 15-nt overhang was synthesized by mixing two completely complementary oligonucleotides (sense-15 and antisense strands in Table S1) in 1×TAE/$Mg^{2+}$ buffer with further incubation for 1 h at 20℃. Next, DNA nanostructures with overhangs were hybridized with the pre-formed siRNA duplex in PBS (137 mM NaCl, 2.7 mM KCl, 10 mM $Na_2HPO_4$, 2 mM $KH_2PO_4$, pH 7.4) at 37℃ for 30 minutes with a final concentration of 100 nM, allowing conjugation of siRNA to the DNA nanostructures. AFM characterization of nanostructure hybridization locus was performed by labeling each nanostructure locus with a biotin at the 5' end of the core strand. Following co-incubation of the biotinylated nanostructure with streptavidin, AFM imaging was performed.

**Preparation of SWCNTs with 15-nt overhang wrapped with DNA $(GT)_{15}$ -15 strand.**

We designed and prepared the $GT_{15}$-15 wrapped SWCNTs where the 15-nt overhangs were complementary to the 15-nt overhangs on the locus of each nanostring monomer, such that the SWCNT and nanostring would hybridize to each other (see detailed sequences in Table S1 and detailed protocols in methods below). To conjugate the SWCNT to the nanostring, the $(GT)_{15}$-15 strand was first dissolved at a concentration of 20 mg/mL in PBS buffer. 1 mg HiPCO SWCNTs was added to 400 µL of this DNA solution, followed by probe-tip sonication with a 3-mm tip at 50% amplitude (~7W) for 20 min in an ice bath. The resulting solution was next centrifuged at 16,100g for 1 h to remove unsuspended SWCNT. Unbound (free) DNA was removed via spin-filtering (Amicon, 100 K) at 1,000g for 6 minutes (5 times) and the concentration of $(GT)_{15}$-15 wrapped SWCNTs was determined with a UV-Vis-nIR spectrometer where SWCNT concentration was calculated in mg/L (absorbance at 632 nm/extinction coefficient of 0.036).

**Conjugation of nanostrings with $(GT)_{15}$-15 SWCNTs.**

Nanostrings with 15-nt hybridization overhangs which were fully complementary with the overhang of $(GT)_{15}$-15 were mixed with the filtered DNA wrapped SWCNTs in 0.5×TAE/$Mg^{2+}$ buffer and incubated at 37℃ for 30 minutes. The conjugate was characterized with AFM by next adding streptavidin to bind to and indicate the biotin labeling position along the nanostring. To better differentiate the nanostring from the SWCNTs, same biotin-specific streptavidin strategy was employed for AFM characterization of the SWCNT-nanostring conjugate. As shown in AFM images in Fig S8, we observe discrete patterns of anchored streptavidin molecules on the surface of SWCNTs, which allowed direct visualization of the successful conjugation.

**Quantitative GFP fluorescence intensity analysis of gene silencing.**

Infiltrated plant leaves were prepared for confocal imaging 3-days post-infiltration with corresponding nanomaterials by cutting a small leaf section of the infiltrated leaf tissue, and inserting the tissue section between a glass slide and cover slip of #1 thickness. 20 µL of water was added between the glass slide and cover slip to keep the leaves hydrated during imaging. A Zeiss LSM 710 confocal microscope was used to image the plant tissue with 488 nm laser excitation and with a GFP filter cube. GFP fluorescence images were obtained at 10x magnification. Confocal imaging data was analyzed to quantify GFP expression across samples. For each sample, 4 biological replicates (4 infiltrations into 4 different plants) were performed, and for each biological replicate, 15 technical replicates (15 non-overlapping confocal field of views from each leaf) were collected. Each field of view was analyzed with custom ImageJ analysis to quantify the GFP fluorescence intensity value for that field of view, and all 15 field of views were then averaged to obtain a mean fluorescence intensity value for that sample. The same protocol was repeated for all 4 biological replicates (4 different plants) per sample, and averaged again for a final fluorescence intensity value, which correlates with the GFP fluorescence intensity of the sample.

**Quantitative PCR (qPCR) experiments and data analysis.**

Two-step qPCR was performed to quantify GFP gene silencing in transgenic mGFP5 *Nicotiana benthamiana* plants with the following commercially-available kits: RNeasy plant mini kit (QIAGEN) for total RNA extraction from leaves, iScript cDNA synthesis kit (Bio-Rad) to reverse transcribe total RNA into cDNA, and PowerUp SYBR green master mix (Applied Biosystems) for qPCR. The target gene in our qPCR was mGFP5 (GFP transgene inserted into *Nicotiana benthamiana*), and EF1 (elongation factor 1) was chosen as the housekeeping (reference) gene. Primers (see detailed sequences in Table S1) for these genes (fGFP, rGFP, fEF1 and rEF1) were ordered from IDT and used without further purification. An annealing temperature of 60˚C was used for qPCR, which was run for 40 cycles. qPCR data was analyzed by the ddCt method to obtain the normalized GFP gene expression-fold change with respect to the EF1 housekeeping gene and control sample. For each sample, qPCR was performed as 3 technical replicates (3 reactions from the same isolated RNA batch), and the entire experiment consisting of independent infiltrations and RNA extractions from different plants was repeated 4 times (4 biological replicates).

**Quantitative Western Blot experiments and data analysis.**

Infiltrated plant leaves were harvested after 72 h and ground in liquid nitrogen to get dry frozen powders. The frozen powders were then transferred to a tube with pre-prepared lysis buffer containing 10 mM Tris/HCl (pH 7.5), 150 mM NaCl, 1 mM EDTA, 0.1% NP-40, 5% glycerol, and 1% cocktail. After lysing at 4℃ overnight, the tube was centrifuged at 10,000 rpm for 20 minutes and the supernatant containing whole proteins was collected to a new tube. After quantification of the total extracted proteins by a Pierce 660 nm Protein Assay (Thermofisher, Prod# 22660), 0.5 µg of normalized total proteins from each sample were analyzed by 12% SDS–PAGE and blotted to a PVDF membrane. The membrane was then blocked for 1 hour using 7.5% BSA in PBST (PBS containing 0.1% Tween20) buffer and rinsed 3 times in PBST buffer, followed by overnight incubation at 4°C with the primary GFP antibody as required (1:2000 dilution, Abcam, ab290). After extensive washing, the corresponding protein bands were probed with a goat anti-rabbit horseradish peroxidase-conjugated antibody (1:5000 dilution, Abcam, ab205718) for 30 min. After 3 washes, the membrane was then developed by incubation with chemiluminescence (Amersham ECL prime kit) plus and imaged by ChemiDoc™ XRS+ System (BIORAD). The intensity of GFP bands were quantified with ImageJ software. To correct for variability in protein expression across different plants and leaves, the GFP extracted from each leaf sample was normalized by the total protein recovered from that leaf tissue.

**Quantitative colocalization analysis of Cy3 labeled nanomaterials with GFP**

Transgenic mGFP5 *Nicotiana benthamiana* plant leaves were infiltrated with 50 µL Cy3-labeled nanostructures to a final nanostructure concentration of 200 nM, and prepared for confocal imaging (12h for DNA materials, and 6h for carbon nanotube related materials) following infiltration. Specifically, one of the single stranded DNA overhangs on the corresponding DNA nanostructure was labeled by Cy3: the H1 strand in HT monomer and nanostring, and one of the four vertices of the tetrahedron (Table S1). SWCNTs were wrapped with Cy3 labeled $GT_{15}$ ssDNA, and the SWCNT-nanostring hybrid was prepared via Cy3 labeling of the nanostring at the nanostring center, which was subsequently conjugated with SWCNT. All three Cy3-labeled structures were infiltrated into mGFP5 *benthamiana* leaves. A small leaf section of the infiltrated leaf tissue was cut, and inserted between a glass slide and cover slip of #1 thickness. 20 µL of water was added between the glass slide and cover slip to keep the leaf sections hydrated during imaging. A Zeiss LSM 710 confocal microscope was used to image the plant tissue with two channels: 488 nm laser excitation with a GFP filter cube and 514 nm laser excitation with a Cy3 filter cube. The images were obtained with air-immersion of the objective at 20x magnification. Confocal imaging data were then analyzed to quantify the colocalization fraction between GFP channel and the Cy3 channel across all samples (image J). For each sample, 4 biological replicates (4 infiltrations into 4 different plants) were performed, and for each biological replicate, 15 technical replicates (15 non-overlapping confocal field of views from each leaf) were collected. Each field of view was analyzed with custom ImageJ analysis software to quantify the colocalization percentage value for that field of view, and all 15 field of views were then averaged to obtain a mean value for that sample. The same protocol was repeated for all 4 biological replicates per sample, and averaged again for a final colocalization value, which correlates with the percent colocalization between the Cy3 and GFP channels for each sample.
